# bMINTY: Enabling Reproducible Management of High-Throughput Sequencing Analysis Results and their Metadata

**DOI:** 10.64898/2026.02.08.704625

**Authors:** Konstantinos Kapelios, Panagiotis Xiropotamos, Haris Manousaki, Charis Sinnis, Vasiliki Kotsira, Theodore Dalamagas, Georgios K. Georgakilas

**Affiliations:** Information Management Systems Institute, ATHENA Research Center, 15125 Marousi, Greece; Department of Computer Science and Biomedical Informatics, University of Thessaly, Lamia, Greece

## Abstract

Due to the large scale of high-throughput sequencing data generation, the community and publishers have established standards for the dissemination of studies that produce and analyze these data. Despite efforts towards Findable, Accessible, Interoperable and Reproducible (FAIR) science, critical obstacles remain. Best practices are not consistently enforced by scientific publishers, and when they are, essential information is fragmented across the methods section, supplementary materials, and public repositories. When attempting to reproduce scientific findings or reuse published data or analyses, researchers often avoid analyzing sequencing data from the ground up. Instead, they prefer to start directly from the post-sequence-alignment information (*e*.*g*., gene expression matrices in transcriptomics). However, existing repositories and workflow-oriented solutions rarely provide a single, portable, queryable resource that integrates this information with the metadata required for downstream reuse.

We introduce bMINTY, a locally deployed web application with an intuitive user interface, for structured management of post-alignment workflow data outputs. bMINTY supports metadata for studies, assays, and analysis assets, including workflows, genome assemblies, genomic intervals, and cell-level entities for single-cell assays. Users may export query results in RO-Crate format, providing machine readable data packages and metadata. To the best of current knowledge, bMINTY is the first framework to bundle all this information in publication-ready, portable packaging designed for reuse. These packages can be included as supplementary material with each publication, accompanied by analysis code deposited in public repositories for downstream *ad hoc* analyses. Together, these practices can promote transparency, efficient reuse of published data, and support FAIR-aligned scientific reproducibility.

## Introduction

### Background and Motivation

Next-generation sequencing (NGS) has transformed genomics over the past decades by enabling high-throughput sequencing of DNA and RNA at steadily decreasing cost^1,2^, marking the beginning of a new era of data-driven molecular biology. Now routinely used in both academia and industry, NGS has been widely adopted through the development of diverse assays spanning major biomedical applications, including molecular biology, metagenomics and genetic disorders among others^3^. Recent advances have further expanded NGS to single-cell and spatially resolved technologies, adding cellular and tissue-context information to molecular profiling^4,5^.

NGS data analysis typically consists of processes that can be grouped into two distinct broad steps: (i) a standard raw sequencing data processing pipeline (SSDPP) and (ii) study-specific downstream analyses^3^. The SSDPP is applied across assays and generally includes quality control, contaminant removal, alignment to a reference genome, detection of enriched regions (*e*.*g*., peak calling in epigenomic profiling assays), and quantification of signal over genomic elements. Its output is typically a two-dimensional matrix with read counts on genes across samples (bulk assays) or cells (single-cell assay). For epigenomic profiling assays, the SSDPP result is usually a collection of signal-enriched genomic regions^6,7^.

In this work, we focus on the SSDPP step, which represents a major bottleneck in NGS data analysis due to its computationally intensive and input/output-heavy nature. After the SSDPP analysis, and to reduce the storage footprint, researchers typically send FASTQ files to cold storage and only keep the results of the final SSDPP step for downstream analyses. In most cases where others would be interested in reusing published data, they would prefer to avoid rerunning the SSDPP steps due to their resource demanding properties, and directly use the SSDPP results.

Despite substantial community efforts (see the *“Related Work and Limitations”* section), the existence of the Findable, Accessible, Interoperable, and Reproducible (FAIR) data principles^8,9^ and standards such as research object crates (RO-Crates)^10^, SSDPP data reuse remains challenging. Assets required to interpret SSDPP outputs, including tools, parameters, reference resources and metadata, are frequently fragmented across manuscripts, supplementary files, public repositories and codebases. As a result, fully reproducing published results or reusing public data with minimal effort remains uncommon.

To address these issues, we designed and developed bMINTY (Figure 1), a lightweight, locally deployable framework that consolidates SSDPP outputs and their essential metadata into a structured, interoperable, and portable resource. Specific contributions of our work include: i) locally organized SSDPP data and metadata with optimized performance, ii) portable and modular architecture based on open-source popular solutions such as SQLite, Django REST application programming interface (API), and ReactJS user interface (UI), iii) assay-agnostic database schema that supports bulk, single-cell, and spatially resolved NGS assays, iv) integration of SSDPP contextual metadata such as study/assay annotations, workflow specifications, genome assemblies, genomic intervals and cells, v) user-friendly capabilities for importing, exploring as well as exporting data, and vi) FAIR-aligned, publication sharing by adopting the RO-Crate standard^10,11^.

**Figure 1.**
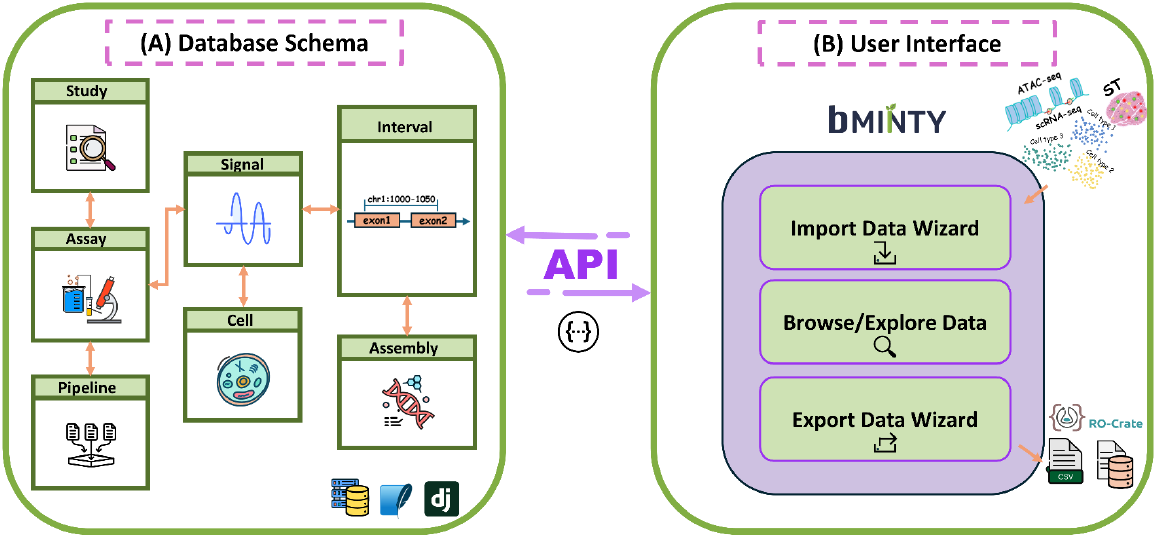
Overview of the bMINTY architecture and core functionality. (A) SQLite- and Django-enabled backend. The database schema has been specifically designed to host metadata and data related to a wide spectrum of high-throughput sequencing assays. Instead of hosting the results of every step of an analytic workflow, we have chosen to focus only on hosting the outcome of the standard raw sequencing data processing pipeline (SSDPP) that starts from the quality control assessment of FASTQ files and ends with counting the amount of overlapping reads with genomic elements (e.g., genes, custom signal-enriched intervals from peak-calling algorithms). The outcome of this workflow is usually a matrix of read counts over genomic elements across one or multiple samples, or BED formatted files as produced by peak calling algorithms. The application programming interface (API) provides a wide spectrum of database querying capabilities that can be used by the bMINTY frontend application or by seamlessly integrating bMINTY with analytic workflows. (B) bMINTY user interface (UI) that offers import/export capabilities and advanced filtering options for querying the information hosted in the database. Data can be uploaded to the database using the import wizard functionality that facilitates the process step-by-step and adheres to the supported hierarchical dependencies of the database structure. The export data wizard supports different types of formats and standards, including RO-Crates.

We envision this portable body of information being the supplementary material of every publication, accompanied by code deposited in public repositories that performs all downstream *ad hoc* analyses, promoting transparency, efficient reuse of published data, and scientific reproducibility.

### Related Work and Limitations

There are numerous published frameworks aiming to improve the *status quo* in FAIR-principled biomedical science, often indexed in bio.tools^12^. To our knowledge, there are no tools available in the literature that aim to address the SSDPP data reuse issue in the way that bMINTY does. However, there are frameworks that attempt to address complementary issues in a FAIR-principled manner. A major category emphasizes reproducible workflows, including Galaxy^13^, Snakemake^14^ and Nextflow^15^. Related approaches extend workflow capabilities with project organization and packaging features, such as PM4NGS^16^, and ReUseData^17^ iRODS^18^ and medna-metadata^19^, while also supporting analytical workflows for genetic and environmental studies. Other solutions, such as openBIS^20^, offer structured data and metadata management for large organizations that perform multi-domain experiments (*e*.*g*., imaging, mass spectrometry, high-throughput sequencing), often requiring tiered authentication/authorization and provenance tracking. Finally, variant-centric tools, including GEMINI^21^, TileDB-VCF^22^, gnomAD DB^23^, VCF-Miner^24^, and GenMasterTable^25^.

Despite these advances, existing frameworks tend to address complementary components of the problem rather than providing an integrated mechanism for day-to-day management and publication-ready reuse of SSDPP outputs. Existing frameworks address important aspects of FAIR-aligned science, but largely in isolation. Solutions oriented around workflows indeed achieve to enhance reproducibility of execution but ensuring findability, structured accessibility, or interoperability of data and metadata associated to the SSDPP is not within their scope. Platforms focusing on metadata improve data annotation and provenance tracking, yet they often rely on centralized systems or complex deployment models that prevent their adaptation to routine day-to-day research operations, while limiting the SSDPP data and metadata portability. Lastly, tools for Variant Call Format (VCF) management and visualization are highly effective for an important but narrow class of genomic assets, albeit lacking support for the broader spectrum of NGS assays. Additionally, at the time of this study, existing solutions do not provide out-of-the-box support for RO-Crates^26^.

Despite the FAIR principles adoption by existing frameworks, the need remains for lightweight, locally deployable solutions that offer structured, interoperable, and reusable organization of the SSDPP results, while fitting seamlessly in the dynamic, fast paced environment of research laboratories. We believe that bMINTY will be an invaluable asset towards this direction.

## Materials and Methods

### bMINTY Implementation

#### Database

The database is stored in a single SQLite file. This format was chosen to enforce data integrity while enabling collaboration and easily sharing the entire system’s state. The database tables are described in Table 1, their fields with descriptions and examples are detailed in Supplementary Table 1, while the database schema is abstracted in Figure 1A (see the GitHub documentation for lengthy details). Briefly, the database’s top level container is the study table. Each study represents a collection of assays with the overarching goal of representing a specific research inquiry. The assay table contains information about the individual experimental protocols performed on samples, such as the tissue of origin and the treatment that was applied prior to the sampling, among others. The study and assay fields have been designed in accordance with the Gene Expression Omnibus (GEO)^27^ metadata structure, to facilitate interoperability with this popular repository. Each assay is linked to a specific SSDPP in the pipeline table, which contains an external repository URL where detailed information about the pipeline has been submitted to (*e*.*g*., WorkflowHub^28^).

**Table 1.** Overview of database tables. The tables are connected in accordance to the hierarchical structure depicted in Figure 1A.

| Database table | Description |
| --- | --- |
| study | Information related to the overarching scientific objective. Each study can be linked to multiple assays. |
| assay | Metadata describing the experimental setup and procedures. Each assay can be linked with a single study, pipeline, and assembly. |
| pipeline | Information regarding the SSDPP used to produce the hosted data. |
| assembly | Information about the genome assembly used in the SSDPP. |
| interval | Metadata related to the genomic intervals exhibiting a signal as measured by the assay. |
| cell | Cell-related information in the case of single-cell assays. |
| signal | The data produced by the SSDPP and information that connects them to the assay, genomic intervals, and cells. |

The genomic data is stored in three discrete tables; interval, signal and cell. Intervals define specific genomic regions along with their metadata, while being linked to an assembly in the assembly table, which stores details regarding the genome assembly version (*e*.*g*., GRCh38) and the public repository (*e*.*g*., Ensembl 115) used in the SSDPP. The signal table records the raw counts or signal enrichments (depending on the assay), acting as a link between them. Finally, the cell table stores metadata for individual cells. In the case of spatially resolved transcriptomics, the coordinates of cells can be stored as well.

#### Backend and Application Programming Interface

bMINTY is designed as a modular application consisting of backend services implemented in the Django framework and a ReactJS UI, operating on a SQLite database format (Figure 1A). Users can optionally deploy both modules as docker containers using a single docker compose command. The Django framework serves as a robust API layer that facilitates rapid development; albeit with data governance, logic management and data validation. Each table of the bMinty database is represented as a separate application in the backend, along with its database properties, test cases, URLs and serializers. In addition, there are separate applications for the database management, such as imports and exports through the persistence layer (SQLite). The REST API can also operate as a standalone service, offering seamless integration in scientific workflows. To facilitate transparency and ease of use, a Swagger UI is provided with the deployment enabling researchers to interactively explore the available endpoints. Shell scripts are provided for data import, rebuilding, monitoring and logging of both frontend and backend containers.

#### bMINTY Import Format, Automated Data Ingestion and Interoperability with Downstream Analyses

For bulk-importing data to the database, users need to adhere to the bMINTY import format (detailed instructions can be found in the GitHub documentation). To facilitate ease-of-use, we have developed a collection of utility scripts, hosted in the GitHub repository, that support the restructuring of frequently used SSDPP results (*e*.*g*., bulk/single-cell/spatial transcriptomics gene expression matrices, narrowPeak or BED) into files formatted according to the import bMINTY standards (Table 2). To facilitate interoperability with downstream analyses, we have also developed utility scripts that receive bMINTY export files and transform them back into the aforementioned popular SSDPP output formats (Table 2). Experienced users can of course adjust the utility scripts based on their own needs, use the API directly or code their own import/output parsers altogether.

**Table 2.** Utility scripts for automated data ingestion and downstream analytics interoperability. The most commonly used formats are supported. Detailed information can be found in the GitHub repository.

| Preparing SSDPP results to be imported to bMINTY |  |
| --- | --- |
| narrowPeak_to_bMINTY.py | Converts narrowPeak formatted files to the bMINTY required format. |
| sc_count_matrix_to_bMINTY.py | Converts single-cell RNA-seq count matrices to the bMINTY required format. |
| spatial_count_matrix_to_bMINTY.py | Converts spatially resolved transcriptomics |
|  | count matrices and spatial coordinates to the bMINTY required format. |
| **Making bMINTY exports interoperable to downstream analyses** |  |
| bMINTY_to_narrowPeak_json.py | Converts bMINTY exported files to narrowPeak format. |
| bMINTY_to_sc_count_matrix_json.py | Converts bMINTY exported files to count matrices with genes as rows and cells as columns. |
| bMINTY_to_spatial_count_matrix_json.py | Converts bMINTY exported files to count matrices with genes as rows and cells as columns, accompanied by a tabular file with spatial coordinates. |
| **API-mediated bulk import** |  |
| bulk_import_from_folder.py | Facilitates data import by allowing users to sequentially ingest SSDPP results via the command-line interface, while generating performance and data metrics. |

**Table 2.** Overview of resource allocation and user interface (UI) responsiveness under varying loads after importing the data and metadata of the 3 use-cases. We measured dataset import duration, average central processing unit and memory usage, as well as the added database entries in total, added signal entries (it is the most impactful table for performance) and database size footprint per dataset. These performance indicators were measured with and without the “omit zero-value signals” option of the data import wizard.

|  | Zero signal included | Zero signal excluded |
| --- | --- | --- |
| **Single-cell RNA-seq (1.46GB folder size)** |  |  |
| **Dataset import duration** | 13m 1s | 1m 14s |
| Added database entries | 63.39M | 2.72M |
| Database size footprint | 3.8GB | 167MB |
| Added signal entries | 63.37M | 2.7M |
| **Spatial transcriptomics (2.2GB folder size)** |  |  |
| Dataset import duration | 18m 57s | 3m 7s |
| Added database entries | 98.39M | 12.85M |
| Database size footprint | 5.6GB | 0.7GB |
| Added signal entries | 98.29M | 12.74M |
| **ATAC-seq (40.43MB folder size)** |  |  |
| Dataset import duration | 6.3s | 6.7s |
| Added database entries | 665.59K | 665.59K |
| Database size footprint | 53MB | 53MB |
| Added signal entries | 332.8K | 332.8K |
| **All datasets** |  |  |
| Database entries | 162.44M | 16.23M |
| Database size | 9.5GB | 0.92GB |
| Avg CPU % (1 thread) | 30.38% | 48.64% |
| Avg RAM | 371MB | 286MB |
| Peak RAM | 834MB | 554MB |
| Signal entries | 162M | 15.78M |

As an alternative to the import process using the UI, bMinty provides a configurable command-line import pipeline, which orchestrates the mounting of local data directories and invokes scripts for updating the database, while computing and storing performance metrics (Table 2). The pipeline can be configured using command-line arguments to adapt to the requirements of each system and research project. For example, the read batch size can be adjusted to better leverage the available deployment resources and, if needed, zero-value signals can be omitted from the import process.

## Results

### Use-Case Data

To demonstrate the applicability of bMINTY, it was used to organize data from three different omics modalities; ATAC-seq, single-cell RNA-seq, and spatial transcriptomics. All datasets and associated metadata were transformed to the bMINTY format using the utility scripts (see the “*bMINTY Import Format, Automated Data Ingestion and Interoperability with Downstream Analyses*” section), prior to upload into the bMINTY database.

The ATAC-seq dataset (GSE100738) was obtained from the ImmGen repository^29^ and consists of 199,785 and 100,986 peaks for replicate 1 (GSM2692171) and 2 (GSM2692172) respectively, derived from Double Negative 1 Thymocytes. Peaks filtered with irreproducible discovery rate (32,021 using an adjusted p-value threshold of 0.05) based on the two replicates were also included. Peaks were imported as custom intervals into the bMINTY database, while the associated signal enrichment values were added to the signal table. This information was complemented with metadata about the study and assay, as well as the genome assembly and workflow used to produce the peak files. In total, this use case included 1 study, 1 assay (GSM2692171 metadata were used as representatives for both replicates and the IDR analysis), 1 pipeline, 1 assembly, ∼332.8K interval, and ∼332.8K signal entries.

The single-cell RNA-seq dataset corresponds to 3,774 cells that were filtered^30^ from the full dataset of 68K human peripheral blood mononuclear cells^31^. Metadata regarding the study and the assay were derived from the corresponding publication. Gene annotations were derived from Ensembl version 82 and populated the genomic interval table of the database, while each expressed gene was associated to each cell’s expression level (raw counts) in the signal table. The uploaded assembly information corresponded to Ensembl version 82, while the workflow information was a dummy entry with minimal information derived from the relevant publication. In total, this use case included 1 study, 1 assay, 1 pipeline, 1 assembly, ∼16.8K interval, ∼3.8K cell, and ∼63.4M signal entries.

For the spatial transcriptomics case, we used the Lung 5-1 sample from the NSCLC dataset from Nanostring, Bruker Spatial Biology^32^. We extracted the available study from the above link and assay metadata from the Giotto object provided by Nanostring, and used those to populate the relevant bMINTY database tables. Gene annotations were derived from GENCODE 49. In total, this use case included 1 study, 1 assay, 1 pipeline, 1 assembly, 980 interval, ∼100K cell, and ∼98.3M signal entries.

### bMINTY User Interface

The frontend is designed using the ReactJS architecture paired with the Material UI library (MUI) for a responsive, dashboard-driven user experience. The UI functionality is fully documented in the GitHub repository with lengthy usage scenarios. The frontend serves as a link between the backend and the database. A new instance of the database can be created by uploading an existing SQLite file or by using an interactive wizard embedded in the bMINTY application (Figure 2A). The wizard provides a step-by-step procedure for importing data from CSV files, following the database schema hierarchical structure. Initially, the user creates a new study or appends to an existing one, followed by a pipeline, an assay and an assembly. Subsequently, the user is prompted to bulk import intervals, cells and signals. To preserve data integrity, an option is provided to remove interval entries that would otherwise result in duplicate database references. In addition, for signals, users can opt to omit zero-value signals to reduce the size of the database and improve query performance, a feature that can be very helpful in the case of single-cell or spatially resolved omics. Finally, the wizard automatically preprocesses the data in vectorized batches, resolves conflicts, and updates indexes and foreign keys to maintain the database references. It then feeds them into a native, highly optimized, SQLite bulk import utility that allows for a single commit transaction of million records without exhausting the system resources.

**Figure 2.**
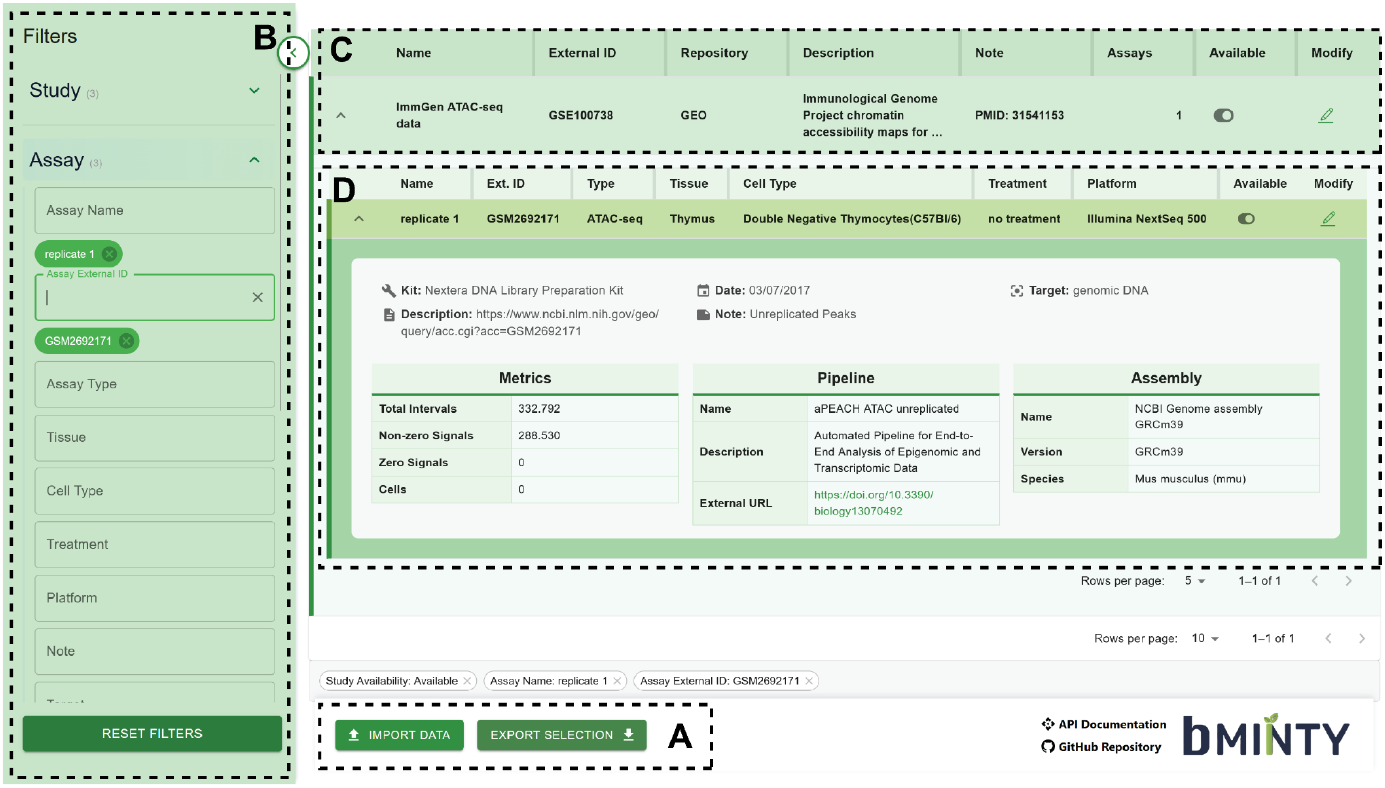
Expanded view of the bMINTY user interface (UI) showcasing the exploration of the imported ATAC-seq, single-cell RNA-seq and spatial transcriptomic datasets. (A) Options that enable the import and export of information to and from the database. The exporting functionality, specifically, allows the exporting of the entire database or only the current filtered view. (B) Options for filtering the information hosted in the database. Users can filter by metadata related to studies, assays, assemblies, intervals and cells. The contents of filter fields are dynamically generated based on the database contents. (C) Top-level view of the database hierarchy showing information about studies. Name and description provide biological context, external id and repository correspond to external sources hosting the original data (if applicable), while the assays field shows the number of assays associated with each study. The note field enables users to store comments related to the study. The availability toggle can be used to hide/show a study and all relevant assays from the UI, while modify allows to alter the contents of a study in the database. (D) Upon expanding a study, all related assays are listed directly below. Information such as name, external id, type, tissue, cell type, treatment, and platform are depicted in the main bar, while the relevant assembly and workflow used to analyze the data as well as metrics/statistics can be viewed after expanding each assay.

Upon uploading data and metadata, users can explore the content by utilizing advanced filtering options to query the database (Figure 2B). The contents of each filter field are dynamically generated based on the content of the corresponding field in the database to avoid engineering static and predefined options, while the main view is automatically reloaded after applying each filter. The filtering results will always follow the hierarchical structure of the database entities. For example, upon the filtering of a specific study (or studies) using the listed name(s), external id(s), repository(ies), description(s) or note(s), only the related assays will be loaded. Subsequently, the users can further filter for a subset of assays based on their names, external ids (if applicable), the related cell types, the tissue origin, treatment scenarios, the platform used to perform the assay, the target molecule, the date that the assay was performed or the kit used to prepare the sample. Lastly, users can further filter the content by interval types and biotypes, assembly names or versions and cell related information.

The main view follows the hierarchical structure of the database table dependencies. At the top level, the user can view information about the studies (Figure 2C). Each study can be expanded to reveal information about the related assays, which can be further expanded to reveal metrics about the datasets, the pipeline used to perform the analysis and the genome assembly that corresponds to the related genomics intervals (Figure 2D). For both studies and assays, users can toggle their visibility through the UI using the “Available” button, and modify the content using the “Modify” button. The “Note” field can be used to store useful notes for future reference, a feature that resembles electronic laboratory notebooks. bMinty also supports exporting the current selection, as defined by the currently applied search filters. Several export formats are available, such as a standalone SQLite file, individual tables in CSV format or a ZIP file containing both. Furthermore, in each of the options mentioned, the export can be encapsulated within a RO-Crate package using the v1.2 specification^11^, which consists of standardized, machine-readable metadata, complying with FAIR data principles.

### bMINTY Performance Under Varying Data Loads

As a proof of concept, bMINTY was used to organize the data derived from three samples corresponding to three distinct types of experiments (Figure 2); ATAC-seq, spatially-resolved transcriptomics and single-cell RNA-seq. bMINTY was deployed as a docker container within a WSL2 environment, running on a system provisioned with 16 GB of RAM and 4 CPUs. The data and metadata were imported using the interactive wizard (Figure 2A), resulting in a SQLite file of 9.5GB (Table 2). This included information about 3 studies, assays, pipelines and assemblies, 350K intervals, 104K cells and 162M signal entries. The importing procedure lasted for approximately 32 minutes. Querying the database using the filtering options provided by the UI (Figure 2B) was not impeded even when hosting such voluminous data sets, while the usage of system resources was kept well within the hardware limits (Table 2). Exporting the database took 6 minutes.

To reduce the size of the database and further improve the performance, we re-imported the spatial transcriptomics and single-cell RNA-seq datasets using the “omit zero-value signals” option. This resulted in 15.78M signal entries instead of the 162M from the initial import, and a database file of 0.92GB. Querying performance was similar to the previous scenario, while exporting the entire database lasted for 2 minutes.

## Discussion

Genomics research has been significantly transformed by the emergence and rapid expansion of NGS technologies^1,2^. As the cost of high-throughput sequencing declined, the community faced critical challenges related to voluminous data management, metadata standardization, transparency regarding the analysis, and most importantly reproducibility of published research outcomes. A prominent response is embodied in the guidelines also known as the FAIR principles^8^.

When publishing a study, it has become standard practice to deposit related sequencing data in public repositories^33–35^. Metadata are typically expected to be deposited in indexing repositories such as GEO^27^, ArrayExpress^36^, and FAIRDOM-SEEK instances^37^. Analysis code is usually shared through well-established collaborative environments and FAIR-principled workflow repositories (*e*.*g*., GitHub, GitLab, WorkflowHub^28^). A practical implementation of the FAIR framework, termed RO-Crates, was recently published and represents a structured archive architecture of all assets related to a research outcome; identifies, provenance, relations, data, and annotations^10^.

Despite these efforts, scientific reproducibility and data reuse still remains an important issue in many fields^8,38,39^ and particularly in both biomedical research^40^ and genomics^41–43^. A recent scoping report from the European Commission^44^ regarding the reproducibility of scientific results highlighted that this issue is driven by cultural, economic, and systemic factors. In 2024, a joint effort between the International Microbiome and Multi-Omics Standards Alliance (microbialstandards.org) and the Genomic Standards Consortium (gensc.org) resulted in a series of community seminars and discussions regarding data reuse and reproducibility within the context of FAIR principles^41^.

From the perspective of a research laboratory, the aforementioned standards and repositories remain too fragmented to be integrated seamlessly into daily operations. In practice, most studies that involve the analysis of NGS data are organized around the SSDPP. Its output (*e*.*g*., count matrices or signal-enriched genomic regions) is the starting point for all downstream, study-specific analyses. Due to the resource-demanding nature of SSDPP execution, researchers typically archive raw FASTQ files in cold storage and only keep the SSDPP outputs for subsequent analyses. Thus, the SSDPP outputs represent the most practically reusable form of NGS data. Every researcher that intends to reuse public NGS data generally prefers to directly use the SSDPP output instead of rerunning the whole workflow.

Interpreting and reusing the SSDPP results is not always straightforward and effortless, since the required contextual information (*e*.*g*., including workflow specifications, software versions, parameter settings, reference genome assemblies, annotation resources, and assay-specific metadata) is frequently spread across multiple sources. Accessing data may require navigating repositories such as GEO^27^. However, the required metadata are often not available at GEO, thus a considerable amount of time has to be dedicated for extracting them from the methods section of the publication or even the supplementary material^41^. The effort of aggregating the information about the SSDPP workflow and any downstream analysis can be even more tedious. The information can be fragmented across the manuscript, supplementary files, and, at best, a public code repository. Consequently, rather than being productive, researchers attempting to reuse SSDPP outputs typically spend a disproportionate amount of time locating, validating, and integrating scattered information. The challenge becomes even greater in multi-omics studies, where data types such as genomics, transcriptomics, proteomics, and spatial transcriptomics must be analyzed together. Taken together, these observations underscore the need for tools that provide structured management of SSDPP outputs together with the metadata required for downstream analyses.

In this context, bMINTY was developed specifically for addressing this issue. Instead of attempting to include information about every possible *ad hoc* genomic analysis, bMINTY focuses on the SSDPP output which is a shared feature between the majority of NGS assays. With the consolidation of metadata, the standard analysis results, workflow descriptions, genome assemblies and annotation resources into a portable database format, bMINTY provides a structured environment that laboratories can adopt without altering their scientific operations or computational preferences. Importantly, this approach embeds reproducibility at the most essential level, before downstream study-specific analyses introduce variation in methodology and interpretation.

The key advantage of bMINTY is that it can be used simultaneously as a tool for organizing daily research, and as an interoperable archival resource supplementing scientific publications. Both frontend and backend can be deployed easily as a single Docker container, thus removing the burden of the sometimes tedious procedure of manually installing dependencies. The programmatic access support through the REST API enables bMINTY to be integrated seamlessly with analytic workflows on *ad hoc* bases. The local web UI ensures ease-of-use and enables researchers to import, browse, filter, and export information from the database efficiently, without compromising data privacy or requiring specialized infrastructure. The portable SQLite packaging can be submitted as supplementary material to published research studies. When exported in a RO-Crate format, combined with code repositories of downstream analyses, it has the potential to promote long term SSDPP data reuse and redefine the concept of reproducible genomics by enabling the full reconstruction of publication results.

Currently, bMINTY can operate only on SQLite databases. SQLite offers flexibility, interoperability and seamless data exchange, however, it lacks scalability. For laboratories that are producing large amounts of data, an efficient database engine (*e*.*g*., PostgreSQL) is required for optimizing query times. Another limitation of bMINTY is that even though the database schema can support storing information for the majority of NGS assays that can associate biological signals to genomic elements, it does not support the storage of variant calling results which is the standard analysis for genetic assays that are very popular and typically performed in bulk for the purposes of genetic studies. There are already well-established tools for storing, organizing, querying, or visualizing VCF content^21–25^, thus this work focuses on addressing the gap in the remaining genomics assay types.

Despite its strengths and limitations, bMINTY does not aim to replace the well-established public repositories, laboratory information management systems, or FAIR-principled standards such as RO-Crates. On the contrary, bMINTY is designed to bridge the dynamic operating environment of individual research laboratories and the rigid, formal requirements for data and metadata cataloguing as set by the community, regulatory bodies, funding agencies and scientific literature publishers. Thus, bMINTY complements instead of competing with existing FAIR-aligned tools, repositories, and standards.

The widespread adoption of frameworks such as bMINTY by the community could play a pivotal role in addressing the reproducibility issues in the field of genomics research. Such tools can lower the effort required to maintain well-structured SSDPP data, metadata and workflow annotations, encouraging RO-Crate-compatible FAIR practices without adding additional overhead at both the data producer as well as the data consumer levels. In the future, the need for principled and interoperable data management systems will only build up, especially with the widespread adoption of multi-omic assays.

## Abbreviations

ATAC-seq: Assay for Transposase-Accessible Chromatin using sequencing
API: Application Programming Interface
ChIP-seq: Chromatin Immunoprecipitation sequencing
CLIP: Cross-Linking Immunoprecipitation
ENA: European Nucleotide Archive
Ensembl: Ensembl Genome Database
FAIR: Findable, Accessible, Interoperable and Reproducible
NGS: Next Generation Sequencing
RNA-seq: RNA Sequencing
REST: Representational State Transfer
RO-Crate: Research Object Crate
SSDPP: standard raw sequencing data processing pipeline
SRA: Sequence Read Archive
SRT: Spatially Resolved Transcriptomics
UI: User Interface
WSL2: Windows Subsystem for Linux 2
BED: Browser Extensible Data
GEO: Gene Expression Omnibus
NSCLC: Non-small-cell lung cancer
VCF: Variant Call Format

## Availability of Source Code and Requirements

Project name: bMINTY

Project homepage: github.com/GeorgakilasLab/bMINTY & https://georgakilaslab.github.io/bMINTY/

Operating system(s): Linux/Windows/MacOS Programming language: Python, ReactJS

Other requirements: Python (version 3.11), Docker (v. 28.2.2) Pandas (v. 2.2.0), Django (v. 5.1.6), Django REST Framework (v. 3.15.2), Django-cors-headers (v. 4.7.0), Django-filter (v. 25.1), ReactJS (v. 19.0.0), React Router DOM (v. 7.3.0), Material UI (MUI) (v. 7.3.7), Axios (v. 1.13.4), ReactFlow (v. 11.11.4), SQLite (v. 3.37.2), roc-validator (v. 0.8.0)

License: GNU General Public License (GPL) version 3 or later.

## Data Availability

The data used in this study are available in Zenodo (https://doi.org/10.5281/zenodo.18516705) as two RO-Crates including bMINTY-compliant SQLite database files. The original ATAC-seq data can be found in GEO with the identifiers GSM2692171 and GSM2692172, the original single-cell RNA-seq data can be downloaded from the 10x Genomics website https://cf.10xgenomics.com/samples/cell/pbmc68k_rds/all_pure_pbmc_data.rds, and the original spatial transcriptomics data can be found at https://nanostring.com/products/cosmx-spatial-molecular-imager/ffpe-dataset/nsclc-ffpe-dataset/.

## Author Contributions

G.K.G. conceptualized the study. K.K. developed the code. G.K.G., V.K., K.K., and T.D. wrote the manuscript. V.K. and P.X. prepared the figures. C.S., P.X. and H.M. prepared the use-case datasets and developed the utility scripts. All authors proofread the manuscript. G.K.G. oversaw the study.

