## Supplementary Text for "bMINTY: Enabling Reproducible Management of High-Throughput Sequencing Analysis Results and their Metadata"

***Supplementary Table 1. Overview of database tables.*** *The tables are connected in accordance to the hierarchical structure depicted in Figure 1A. The table summarizes the database table fields with descriptions and examples, as well as the types of relationship between them. The examples shown correspond to the ATAC-Seq use-case. The * signifies mandatory fields and the ** fields that are automatically set by the bMINTY backend.*

| Column name | Description | Example |
|---|---|---|
| **study** | | |
| study_id** | primary key | 1 |
| external_id* | unique study id that can be used to find the study in an external repository | GSE100738 |
| external_repo | name of the external repository that the external_id corresponds to | GEO |
| name* | a short name that describes the study | ImmGen ATAC-seq data |
| description | a lengthy description of the study | Immunological Genome Project chromatin accessibility maps for 86 different immunocytes (ATAC-seq). Immune cell populations were isolated in high-purity by flow cytometry. |
| availability | a boolean flag that toggles the visibility of the study in the UI | 1 |
| **pipeline** | | |
| pipeline_id** | primary key | 1 |
| name* | a short name that describes the pipeline | aPEACH ATAC unreplicated |

| | | |
|---|---|---|
| description | a lengthy description of the pipeline | Automated Pipeline for End-to-End Analysis of Epigenomic and Transcriptomic Data |
| external_url* | a url of an external repository that the pipeline has been submitted to | https://doi.org/10.3390/biology13070492 |
| **assembly** | | |
| assembly_id** | primary key | 1 |
| name* | name of the external repository that hosts the genome assembly | NCBI Genome assembly GRCm39 |
| version* | commonly used name of the genome assembly version | GRCm39 |
| species | the 3-letter nomenclature of the species that the assembly corresponds to | Mus musculus (mmu) |
| **assay** | | |
| assay_id** | primary key | 1 |
| external_id* | unique assay id that can be used to find the study in an external repository | GSM2692171 |
| type* | type of the assay | ATAC-seq |
| target | target molecule of the assay (protocol-dependent) | genomic DNA |
| name* | a short name that uniquely describes the assay | replicate 1 |
| tissue | the tissue origin of the sample that was used to perform the assay | Thymus |
| cell_type | the cell type that was isolated from the sample to perform the assay (if applicable) | Double Negative Thymocytes(C57Bl/6) |
| treatment* | a very short description of the treatment that was applied to the biological system prior to taking the sample and performing the assay | not applicable |
| date | date at which the assay was performed | 03/07/2017 |
| platform* | the platform that was used to perform the assay | Illumina NextSeq 500 |
| kit | the kit that was used to prepare the sample that was used to perform the assay | Nextera DNA Library Preparation Kit |
| description | a lengthy description of the assay | https://www.ncbi.nlm.nih.gov/geo/query/acc.cgi?acc=GSM2692171 |
| availability | a boolean flag that toggles the visibility of | 1 |

| | | |
|---|---|---|
| | the assay in the UI | |
| study_id** | foreign key (study) | 1 |
| pipeline_id** | foreign key (pipeline) | 1 |
| interval_count** | total genomic intervals associated with the assay (calculated once when the interval table is populated during import) | 332792 |
| non_zero_cells** | total cells with non-zero signal associated with the assay (calculated once when the cell table is populated during import, only applicable to single-cell assays) | 0 |
| zero_cells** | total cells with zero signal associated with the assay (calculated once when the cell table is populated during import, only applicable to single-cell assays) | 0 |
| **interval** | | |
| interval_id** | primary key | 1 |
| external_id | unique genomic interval id (see documentation for more details), set by the user | 32413 |
| parental_id | the external_id of a parental genomic interval to support hierarchical relationships | null |
| name | name of the genomic interval as assigned by an external repository (in the case of genes) or the user (in the case of custom intervals) | preT.DN1_peak_1 |
| type* | type of the interval as assigned by an external repository (in the case of genes) or the user (in the case of custom intervals) | ATAC_peaks |
| biotype | biotype of the genomic interval as assigned by an external repository | |
| chromosome* | chromosome from which the genomic interval originates | GL456210.1 |
| start* | chromosomal start coordinate of the genomic interval | 159375 |
| end* | chromosomal end coordinate of the genomic interval | 159581 |
| strand* | DNA strand on which the genomic interval is located | + |
| summit | chromosomal coordinate of the genomic interval summit (only applicable to peak-like intervals) | 23 |

| | | |
|---|---|---|
| assembly_id** | foreign key (assembly) | 1 |
| **cell** | | |
| cell_id** | primary key | null |
| name* | name of the cell that is typically given by analytic pipelines in the form of a DNA barcode | null |
| type* | resolution type reflecting the capture technology | null |
| label | cell-type that the cell corresponds too (if applicable) | null |
| x_coordinate | x coordinate of the cell relative to the field of view (applicable to spatial technologies) | null |
| y_coordinate | y coordinate of the cell relative to the field of view (applicable to spatial technologies) | null |
| z_coordinate | z coordinate of the cell relative to the field of view (applicable to spatial technologies) | null |
| assay_id** | foreign key (assay) | null |
| **signal** | | |
| signal_id** | primary key | 507 |
| signal* | experimental signal measured overlapping a genomic interval in the form of raw or normalized read counts | 357.195 |
| p_value | level of statistical significance for the measured signal (only applicable for certain protocols) | 421.397 |
| padj_value | adjusted level of statistical significance for the measured signal (only applicable for certain protocols) | 197.117 |
| assay_id** | foreign key (assay) | 1 |
| interval_id** | foreign key (interval) | 1 |
| cell_id** | foreign key (cell) | null |
